# Assortative mating on ancestry-variant traits in admixed Latin American populations

**DOI:** 10.1101/177634

**Authors:** Emily T. Norris, Lavanya Rishishwar, Lu Wang, Andrew B. Conley, Aroon T. Chande, Adam M. Dabrowski, Augusto Valderrama-Aguirre, I. King Jordan

**Author notes:** Corresponding author: 950 Atlantic Drive, Atlanta, GA 30332.

## Abstract

**Background:** Assortative mating is a universal feature of human societies, and individuals from ethnically diverse populations are known to mate assortatively based on similarities in genetic ancestry. However, little is currently known regarding the exact phenotypic cues, or their underlying genetic architecture, which inform ancestry-based assortative mating.

**Results:** We developed a novel approach, using genome-wide analysis of ancestry-specific haplotypes, to evaluate ancestry-based assortative mating on traits whose expression varies among the three continental population groups – African, European, and Native American – that admixed to form modern Latin American populations. Application of this method to genome sequences sampled from Colombia, Mexico, Peru, and Puerto Rico revealed widespread ancestry-based assortative mating. We discovered a number of anthropometric traits (body mass, height, facial development and waist-hip ratio) and neurological attributes (educational attainment and schizophrenia) that serve as phenotypic cues for ancestry-based assortative mating. Major histocompatibility complex (MHC) loci show population-specific patterns of both assortative and disassortative mating in Latin America. Ancestry-based assortative mating in the populations analyzed here appears to be driven primarily by African ancestry.

**Conclusions:** This study serves as an example of how population genomic analyses can yield novel insights into human behavior.

## Background

Mate choice is a fundamental dimension of human behavior with important implications for population genetic structure and evolution [1-3]. It is widely known that humans choose to mate assortatively rather than randomly. That is to say that humans, for the most part, tend to choose mates that are more similar to themselves than can be expected by chance. Historically, assortative mating was based largely on geography, whereby partners were chosen from a limited set of physically proximal individuals [4]. Over millennia, assortative mating within groups of geographically confined individuals contributed to genetic divergence between groups, and the establishment of distinct human populations, such as the major continental population groups recognized today [5-7].

However, the process of geographic isolation followed by population divergence that characterized human evolution has not been strictly linear. Ongoing human migrations have continuously brought previously isolated populations into contact; when this occurs, the potential exists for once isolated populations to admix, thereby forming novel population groups [8]. Perhaps the most precipitous example of this process occurred in the Americas, starting just over 500 years ago with the arrival of Columbus in the New World [9]. This major historical event quickly led to the co-localization of African, European and Native American populations that had been (mostly) physically isolated for tens of thousands of years [10]. As can be expected, the geographic reunification of these populations was accompanied, to some extent, by genetic admixture and the resulting formation of novel populations. This is particularly true for populations in Latin America, which often show high levels of three-way genetic admixture between continental population groups [11-15].

Nevertheless, modern admixed populations are still very much characterized by non-random assortative mating. Assortative mating in modern populations has been shown to rest on a variety of traits, including physical (stature and pigmentation) and neurological (cognition and personality) attributes. For example, numerous studies have demonstrated an influence of similarities in height and body mass on mate choice [2, 16-18]. In addition, assortative mating has been observed for diverse neurological traits, such as educational attainment, introversion/extroversion and even neurotic tendencies [19-24]. Harder to classify traits related to personal achievement (income and occupational status) and culture (values and political leanings) also impact patterns of assortative mating [19, 25, 26]. Odor is one of the more interesting traits implicated in mate choice, and it has been linked to so-called disassortative (or negative assortative) mating, whereby less similar mates are preferred. Odor-based disassortative mating has been attributed to differences in genes of the major histocompatibility (MHC) locus, which functions in the immune system, based on the idea that combinations of divergent human leukocyte antigen (HLA) alleles provide a selective advantage via elevated host resistance to pathogens [27, 28].

Ancestry is a particularly important determinant of assortative mating in modern admixed populations [29, 30]. Studies have shown that individuals in admixed Latin American populations tend to mate with partners that have similar ancestry profiles. For example, partners from both Mexican and Puerto Rican populations have significantly higher ancestry similarities than expected by chance [24, 31]. In addition, a number of traits that have been independently linked to assortative mating show ancestry-specific differences in their expression [32]. Accordingly, ancestry-based mate choice has recently been related to a limited number of physical (facial development) and immune-related (MHC loci) traits [24].

The studies that have uncovered the role of genetic ancestry in assortative mating among Latinos have relied on estimates of global ancestry fractions between mate pairs [24, 31]. Given the recent accumulation of numerous whole genome sequences from admixed Latin American populations – along with genome sequences from global reference populations [7] – it is now possible to characterize local genetic ancestry for individuals from admixed American populations [12, 33, 34]. In other words, the ancestral origins for specific chromosomal regions (haplotypes) can be assigned with high confidence for admixed individuals [35]. For the first time here, we sought to evaluate the impact of local ancestry on assortative mating in admixed Latin American populations. Since the genetic variants that influence numerous phenotypes have been mapped to specific genomic regions, we reasoned that a focus on local ancestry could help to reveal the specific phenotypic drivers of ancestry-based assortative mating.

Our approach to this question entailed an integrated analysis of local genetic ancestry and the genetic architecture of a variety of human traits thought to be related to assortative mating. Assortative mating results in an excess of homozygosity, whereas disassortative mating yields excess heterozygosity. It follows that assortative (or disassortative) mating based on local ancestry would yield an excess (or deficit) of ancestry homozygosity at specific genetic loci. In other words, for a given population, a locus implicated in ancestry-based assortative mating would be more likely to have the same ancestry at both pairs of haploid chromosomes within individuals than expected by chance. We developed a test statistic – the assortative mating index (AMI) – that evaluates this prediction for individual gene loci, and we applied it to sets of genes that function together to encode polygenic phenotypes. We find evidence of substantial local ancestry-based assortative mating, and far less disassortative mating, for four admixed Latin American populations across a variety of anthropometric, neurological and immune-related phenotypes. Our approach also allowed us to assess the specific ancestry components that drive patterns of assortative and disassortative mating in these populations.

## Results

### Global and local genetic ancestry in Latin America

We compared whole genome sequences from four admixed Latin American populations, characterized as part of the 1000 Genomes Project (1KGP) [7] to genome sequences and whole genome genotypes from a panel of 34 global reference populations from Africa, Europe and the Americas (Table 1 and Additional file 1: Figure S1). The program ADMIXTURE [39] was used to infer the continental genetic ancestry fractions – African, European and Native American – for individuals from the four Latin American populations (Additional file 1: Figure S2). Distributions of individuals’ continental ancestry fractions illustrate the distinct ancestry profiles of the four populations (Fig. 1). Puerto Rico and Colombia and show the highest European ancestry fractions along with the highest levels of three-way admixture. These two populations also have the highest African ancestry fractions; although, all four populations have relatively small fractions of African ancestry. Peru and Mexico show more exclusively Native American and European admixture, with Peru having by far the largest Native American ancestry fraction.

**Table 1.** Human populations analyzed in this study. Populations are organized into continental groups, for both ancestral and admixed Latin American populations, and the number of individuals in each population and group is shown.

|  | Dataset<sup>1</sup> | Geographical Source | Short | *n* |  | Dataset<sup>1</sup> | Geographical Source | *n* |
| --- | --- | --- | --- | --- | --- | --- | --- | --- |
| Africa (*n*=547) | 1KGP | Esan in Nigeria | ESN | 99 | Native American (*n*=280) | HGDP | Pima in Mexico | 14 |
|  | 1KGP | Gambian in Western Division, The Gambia | GWD | 113 |  | HGDP | Maya in Mexico | 21 |
|  | 1KGP | Luhya in Webuye, Kenya | LWK | 99 |  | Reich et al | Tepehuano in Mexico | 25 |
|  | 1KGP | Mende in Sierra Leone | MSL | 85 |  | Reich et al | Mixtec in Mexico | 5 |
|  | 1KGP | Yoruba in Ibadan, Nigeria | YRI | 108 |  | Reich et al | Mixe in Mexico | 17 |
|  | HGDP | Mandenka |  | 22 |  | Reich et al | Zapotec in Mexico | 43 |
|  | HGDP | Yoruba |  | 21 |  | Reich et al | Kaqchikel in Guatemala | 13 |
| Europe (*n*=471) | 1KGP | Finnish in Finland | FIN | 99 |  | Reich et al | Kogi in Colombia | 4 |
|  | 1KGP | British in England & Scotland | GBR | 90 |  | Reich et al | Waunana in Colombia | 3 |
|  | 1KGP | Iberian populations in Spain | IBS | 107 |  | Reich et al | Embera in Colombia | 5 |
|  | 1KGP | Toscani in Italy | TSI | 107 |  | Reich et al | Guahibo in Colombia | 6 |
|  | HGDP | Russian |  | 25 |  | Reich et al | Piapoco in Colombia | 7 |
|  | HGDP | Orcadian |  | 15 |  | Reich et al | Inga in Colombia | 9 |
|  | HGDP | French |  | 28 |  | Reich et al | Wayuu in Colombia | 11 |
| Admixed (*n*=347) | 1KGP | Colombian in Medellin, Colombia | CLM | 94 |  | HGDP | Karitiana in Brazil | 14 |
|  | 1KGP | Peruvian in Lima, Peru | PEL | 85 |  | HGDP | Suruí in Brazil | 8 |
|  | 1KGP | Mexican Ancestry in LA, California | MXL | 64 |  | Reich et al | Ticuna in Brazil | 6 |
|  | 1KGP | Puerto Rican in Puerto Rico | PUR | 104 |  | Reich et al | Quechua in Peru | 40 |
|  |  |  |  |  |  | Reich et al | Aymara in Bolivia | 23 |
|  |  |  |  |  |  | Reich et al | Guarani in Paraguay | 6 |

<sup>1</sup> 1KG = 1000 Genomes Project; HGDP = Human Genome Diversity Panel; Reich et al<sup>47</sup>

**Figure 1.**
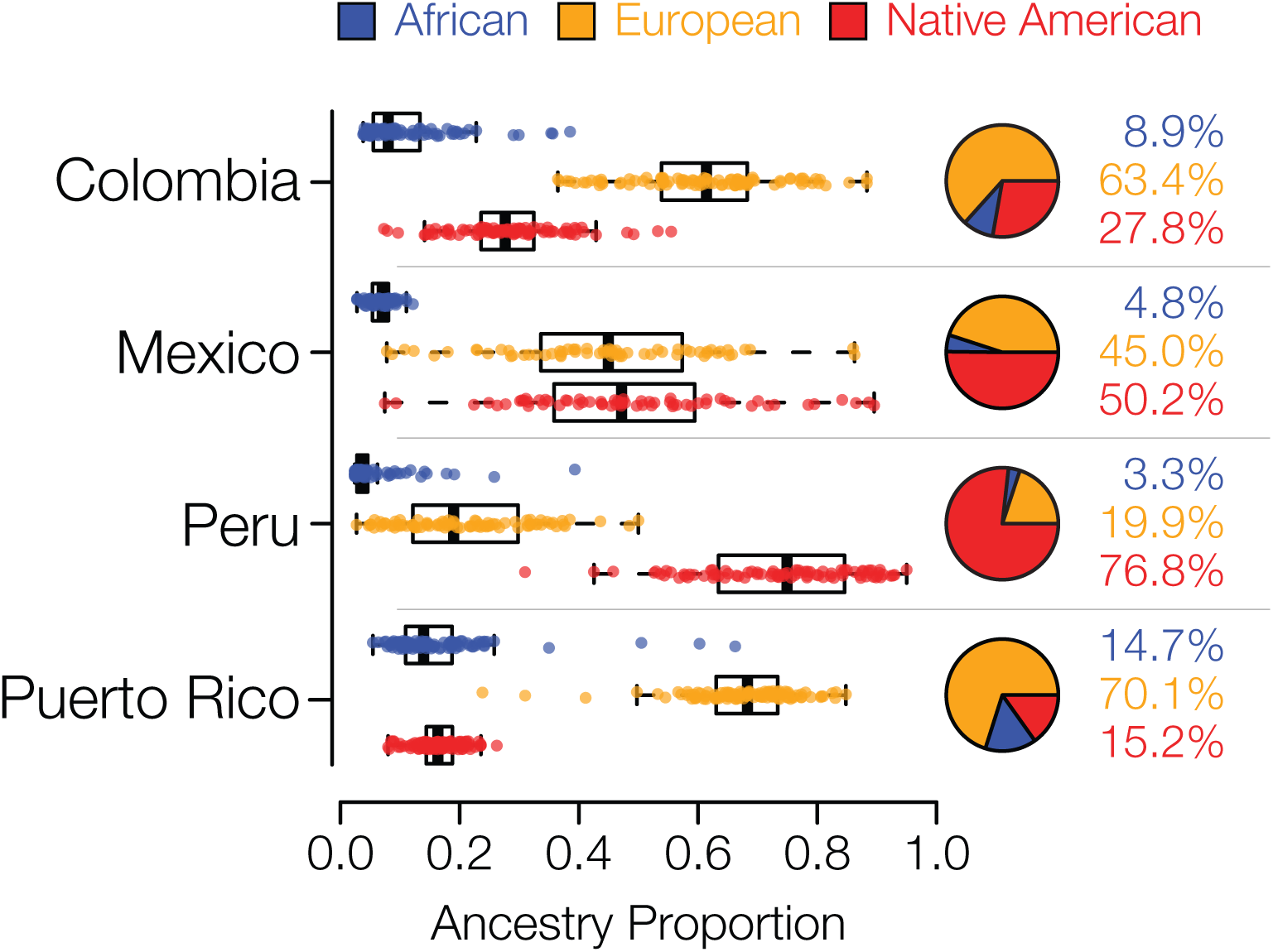
Genetic ancestry proportions for the admixed Latin American populations studied here. For each population, distributions and average values are shown for African (blue), European (orange) and Native American (red) ancestry.

The program RFMix [35] was used to infer local African, European and Native American genetic ancestry for individuals from the four admixed Latin American populations analyzed here. RFMix uses global reference populations to perform chromosome painting, whereby the ancestral origins of specific haplotypes are characterized across the entire genome for admixed individuals. Only haplotypes with high confidence ancestry assignments (≥99%) were taken for subsequent analysis. Examples of local ancestry assignment chromosome paintings for representative admixed individuals from each population are shown in Additional file 1: Figure S3. The overall continental ancestry fractions for admixed genomes calculated by global and local ancestry analysis are highly correlated, and in fact virtually identical, across all individuals analyzed here, in support of the reliability of these approaches to ancestry assignment (Additional file 1: Figure S4).

### Assortative mating and local ancestry in Latin America

We analyzed genome-wide patterns of local ancestry assignment in order to assess the evidence for assortative mating based on local ancestry in Latin America (Fig. 2a). For each individual, the ancestry assignments for pairs of haplotypes at any given gene were evaluated for homozygosity (*i.e.*, the same ancestry on both haplotypes) or heterozygosity (*i.e.*, different ancestry on both haplotypes) (Fig 2b). For each gene, across all four populations, the observed values of ancestry homozygosity and heterozygosity were compared to the expected values in order to compute gene- and population-specific assortative mating index (AMI) values. AMI is computed as a log odds ratio as described in the Methods. The expected values of local ancestry homozygosity and heterozygosity used for the AMI calculations are based on a Hardy-Weinberg triallelic model with the three allele frequencies computed as the locus-specific ancestry fractions. High positive AMI values result from an excess of observed local ancestry homozygosity and are thereby taken to indicate assortative mating based on shared local genetic ancestry. Conversely, low negative AMI values indicate excess local ancestry heterozygosity and disassortative mating.

**Figure 2.**
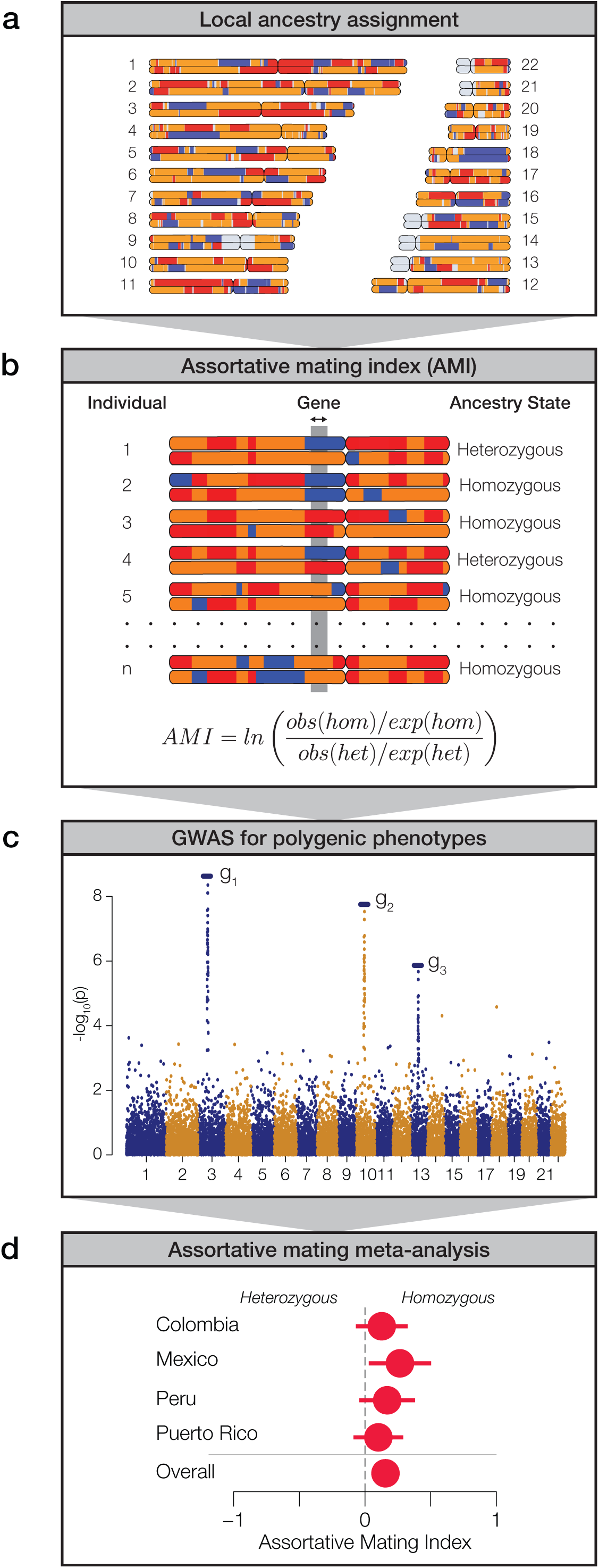
Approach used to measure assortative mating on local ancestry. **(a)** Local ancestry is assigned for specific haplotypes across the genome: African (blue), European (orange), and Native American (red). **(b)** Within individual genomes, genes are characterized as homozygous or heterozygous for local ancestry. For any given population, at each gene locus, the assortative mating index (AMI) is computed from the observed and expected counts of homozygous and heterozygous gene pairs. **(c)** Data from genome-wide association studies (GWAS) are used to evaluate polygenic phenotypes. **(d)** Meta-analysis of AMI values is used to evaluate the significance of ancestry-based assortative mating for polygenic phenotypes.

While we were interested in exploring the relationship between local genetic ancestry and assortative mating, we recognized that mate choice is based on phenotypes rather than genotypes *per se*. Since phenotypes are typically encoded by multiple genes, expressed in the context of their environment, we used data from genome-wide association studies (GWAS) to identify sets of genes that function together to encode polygenic phenotypes (Fig. 2c). We combined data from several GWAS database sources in order to curate a collection of 106 gene sets that have been linked to the polygenic genetic architecture of a variety of human traits. These gene sets range in size from 2 to 212 genes and include a total of 986 unique genes (Additional file 1: Figure S5). We focused on phenotypes that are known or expected to influence mate choice and thereby impact assortative mating patterns. These phenotypes fall into three broad categories: anthropometric traits (*e.g.*, body shape, stature and pigmentation), neurological traits (*e.g.*, cognition, personality and addiction) and immune response (HLA genes). Finally, we used a meta-analysis of the AMI values for the sets of genes that underlie each polygenic phenotype in order to evaluate the impact of local ancestry on assortative mating (Fig. 2d).

We compared the distributions of observed versus expected AMI values to assess the overall evidence for local ancestry-based assortative mating in Latin America. Expected AMI values were computed via permutation analysis by randomly combining pairs of haplotypes into diploid individuals in order to approximate random mating. The distribution of the expected AMI values is narrow and centered around 0, whereas the observed AMI values have a far broader distribution and tend to be positive (expected AMI μ=-0.01, σ=0.03, observed AMI μ=0.11, σ=0.14; Fig. 3a). When all four admixed Latin American populations are considered together, the mean observed AMI value is significantly greater than the expected mean AMI (*t*=18.14, *P*=8.12e-56). The same trend can be seen when all four populations are considered separately (Additional file 1: Figure S6). Mean observed AMI values vary substantially across populations, with Mexico showing the highest levels of local ancestry-based assortative mating and Puerto Rico showing the lowest (Fig. 3b). There is also substantial variation seen for the extent of assortative mating among the three broad functional categories of phenotypes (Fig. 3c). Local ancestry-based assortative mating is particularly variable for HLA genes, with high levels of assortative mating seen for Mexico and evidence for disassortative mating seen for Colombia and Puerto Rico. Anthropometric traits tend to show higher levels of local ancestry-based assortative mating across all four populations compared to neurological traits.

**Figure 3.**
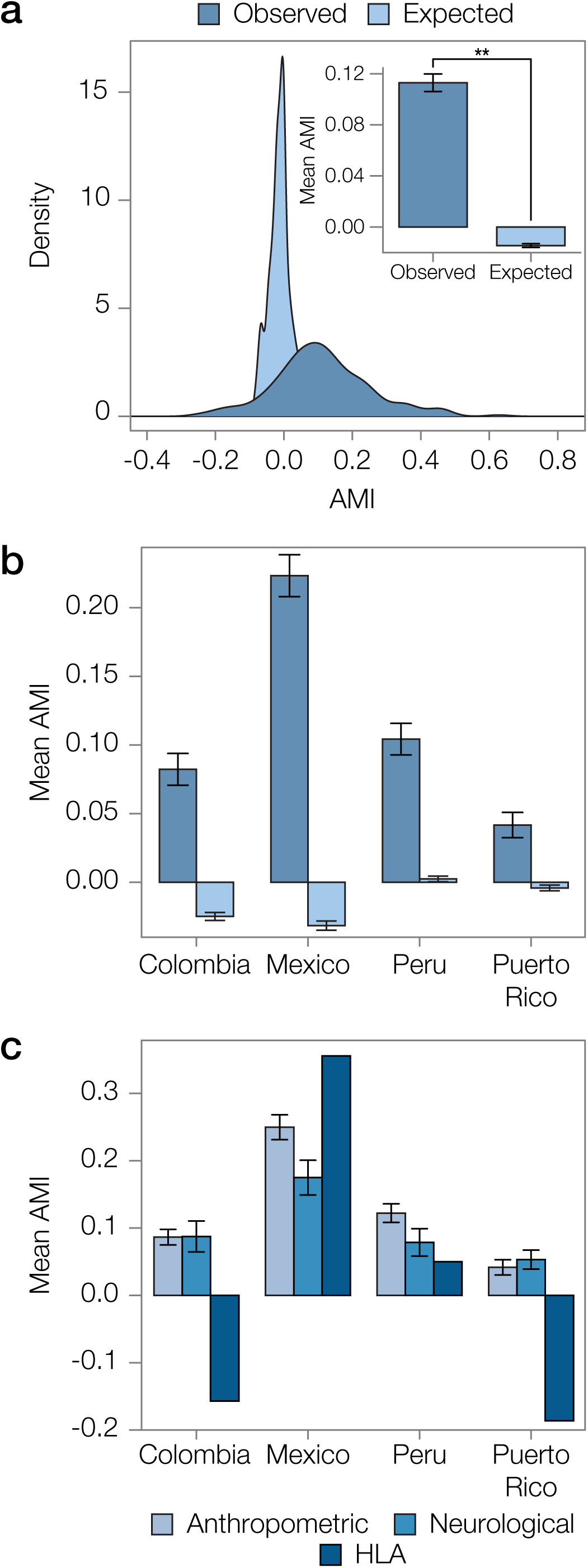
Overview of ancestry-based assortative mating in the four admixed Latin American populations analyzed here. **(a)** Distributions of observed and expected AMI values for all four populations. Inset: Mean observed and expected AMI values (±se) for all four populations. Significance between mean observed and expected AMI values (P=8.12e-56) is indicated by two asterisks. **(b)** Observed and expected average AMI values (±se) across all polygenic phenotype gene sets are shown for each population. **(c)** Average AMI values (±se) for each population are shown for the three main phenotype functional categories characterized here: anthropometric, neurological, and human leukocyte antigen (HLA) genes.

In addition to the permutation test that we used to compute expected AMI values based on randomly paired haplotypes, we also performed a simulation analysis using a population genetic model of assortative mating in order to validate the performance of the AMI test statistic (Additional file 1: Figure S7). We were particularly interested in exploring the potential effects of different ancestry proportions among the populations analyzed here, and different gene set sizes, on computed AMI values. The population genetic model that we used to simulate assortative mating combines Hardy-Weinberg genotype expectations with a single parameter α that represents the fraction of the population that mates assortatively. Details of how this model was implemented to simulate AMI values for the four populations can be found in the Methods section. The population genetic simulation shows that our AMI test statistic is fairly sensitive to low values of the assortative mating parameter α. We also show that AMI values are not biased in any particular direction based on the overall ancestry fractions observed for each population. For example, according to the simulation, Colombia should have the highest overall AMI values, followed by Puerto Rico, Mexico and Peru. This order is completely different from what is seen for the observed AMI values, where Mexico shows the highest mean value, followed by Peru, Colombia and Puerto Rico (Fig. 3b). The population genetic simulation does show that the size of the gene set being analyzed influences the sensitivity of the AMI test statistic. Larger gene sets show greater evidence for assortative mating at the same α parameter values compared with smaller gene sets.

### Local ancestry-based assortative mating for polygenic phenotypes

When considered together, observed AMI levels are enriched for positive values compared to the expected values based on randomly paired haplotypes, indicative of an overall trend of assortative mating based on local ancestry in admixed Latin American populations (Fig. 3a and Additional file 1: Figure S6). We evaluated polygenic phenotypes individually to look for the strongest examples of traits linked to local ancestry-based assortative mating and to evaluate traits that show either similar or variable assortative mating trends across populations. We computed AMI values for 106 polygenic phenotypes across the four populations; the expected and observed AMI values for all traits are shown in Additional file 1: Figure S8. As can be seen for the overall patterns of assortative mating, individual polygenic phenotypes show more extreme positive (for most cases) and negative (in a few cases) AMI values in the four admixed Latin American populations than can be expected for randomly mating populations.

There are 15 polygenic phenotypes that have statistically significant AMI values, after correction for multiple tests, indicative of local ancestry-based assortative mating (*q*<0.05; Fig. 4a). The majority of the statistically significant cases of assortative mating are seen in the Mexican population (8 out of 15), and the anthropometric functional category is most commonly seen among the significant phenotypes (12 out of 15). Height is the most commonly observed phenotype among the significant cases, appearing 6 times in three out of the four populations analyzed here (Colombia, Mexico and Peru). Body mass index is the next most common phenotype, with four significant cases in two populations (Mexico and Peru). The only neurological traits that show significant evidence of assortative mating are schizophrenia (Mexico and Peru) and educational attainment (Mexico). Puerto Rico was the only population that did not show any individual phenotypes with significant evidence of assortative mating, consistent with its low overall AMI values (Fig. 3b and Additional file 1: Figure S6). A list of these significant traits, including references to the literature where the trait single nucleotide polymorphism (SNP)-associations were originally reported, is provided in Additional file 1: Table S1.

**Figure 4.**
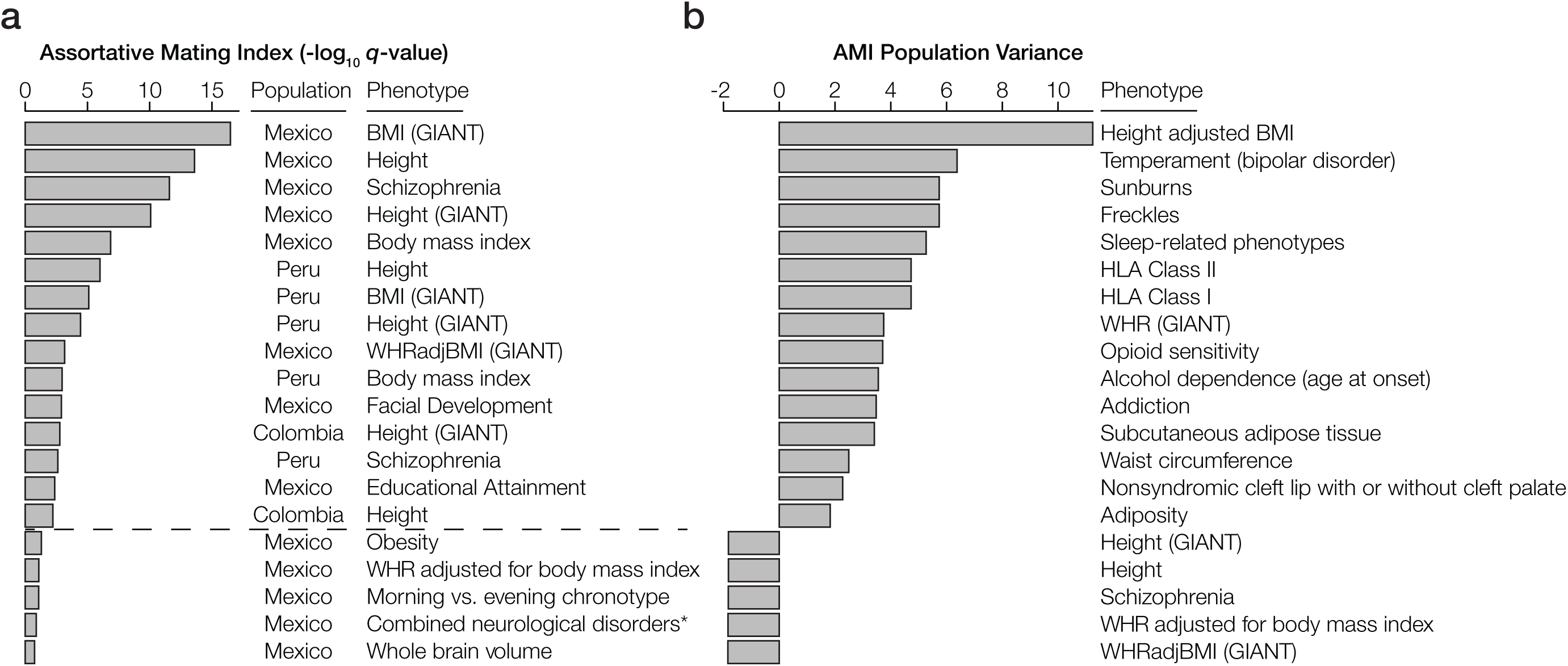
Phenotypes with statistically significant patterns of assortative mating within and among populations. **(a)** The top 20 phenotypes with the highest, and most statistically significant, assortative mating values (AMI) seen within any individual population. All AMI values shown are significant at P<0.05, and the dashed line corresponds to a false discovery rate q-value cutoff of 0.05. **(b)** The top 20 phenotypes with the highest or lowest, and most statistically significant, AMI variance levels across populations. Across population variance levels are normalized using the average AMI population variance level for all phenotypes. All AMI variance levels shown are significant at q<0.05. The highest variance (most dissimilar patterns) of the AMI are at the top, while the lowest variance (most similar patterns) of AMI are at the bottom.

In addition to evaluating individual phenotypes for statistically significant AMI values, we also looked for polygenic phenotypes that showed the most similar or dissimilar patterns of assortative mating across the four admixed Latin American populations. The top 20 phenotypes with the highest and lowest population variance are shown in Fig. 4b (all are statistically significant at *q*<0.05). The polygenic phenotypes with the most variance in population-specific AMI values show more functional diversity compared to the phenotypes with the strongest signals for assortative mating. All three functional categories are represented among the highly population variant phenotypes, and the highly variant phenotypes consist of both assortative and disassortative mating cases (specifically the HLA genes that are described in more detail below). Neurological phenotypes are particularly enriched among the variant cases, including temperament and several addiction-related phenotypes: opioid sensitivity, alcohol dependence and general addiction. Interestingly, all of the least variant phenotypes – height, waist-hip ratio and schizophrenia – are also found among the most significant cases of assortative mating, attesting to a pervasive role in ancestry-based assortative mating for these traits. A list of the population (in)variant traits, including references to the literature where the trait SNP-associations were originally reported, is provided in Additional file 1: Table S1.

Given the evidence of significant local ancestry-based assortative mating that we observed for a number of traits, we evaluated whether there were particular ancestry components that were most relevant to mate choice. In other words, we asked whether the excess counts of observed ancestry homozygosity or heterozygosity are linked to specific local ancestry assignments: African, European and/or Native American. For significant polygenic phenotype gene sets of interest, we computed the observed versus expected ancestry homozygosity for each ancestry separately across all genes in the set (Fig. 5). Height is an anthropometric trait for which Colombia, Mexico and Peru show significant evidence of assortative mating after correction for multiple tests (*q*<0.05; Fig 5a), and Puerto Rico shows nominally significant assortative mating for this same trait (*P*<0.05). In Colombia, Peru and Puerto Rico, assortative mating for this polygenic phenotype is driven by an excess of African homozygosity, whereas in Mexico there is a lack of African homozygosity. The neurological disease schizophrenia shows statistically significant assortative mating in Mexico and Peru (*q*<0.05), with marginally significant values in Colombia and Puerto Rico (Fig. 5b). Patterns of assortative mating for this trait in Mexico and Peru are driven mainly by European ancestry, whereas Colombia and Puerto Rico show an excess of African ancestry homozygosity for this same trait.

**Figure 5.**
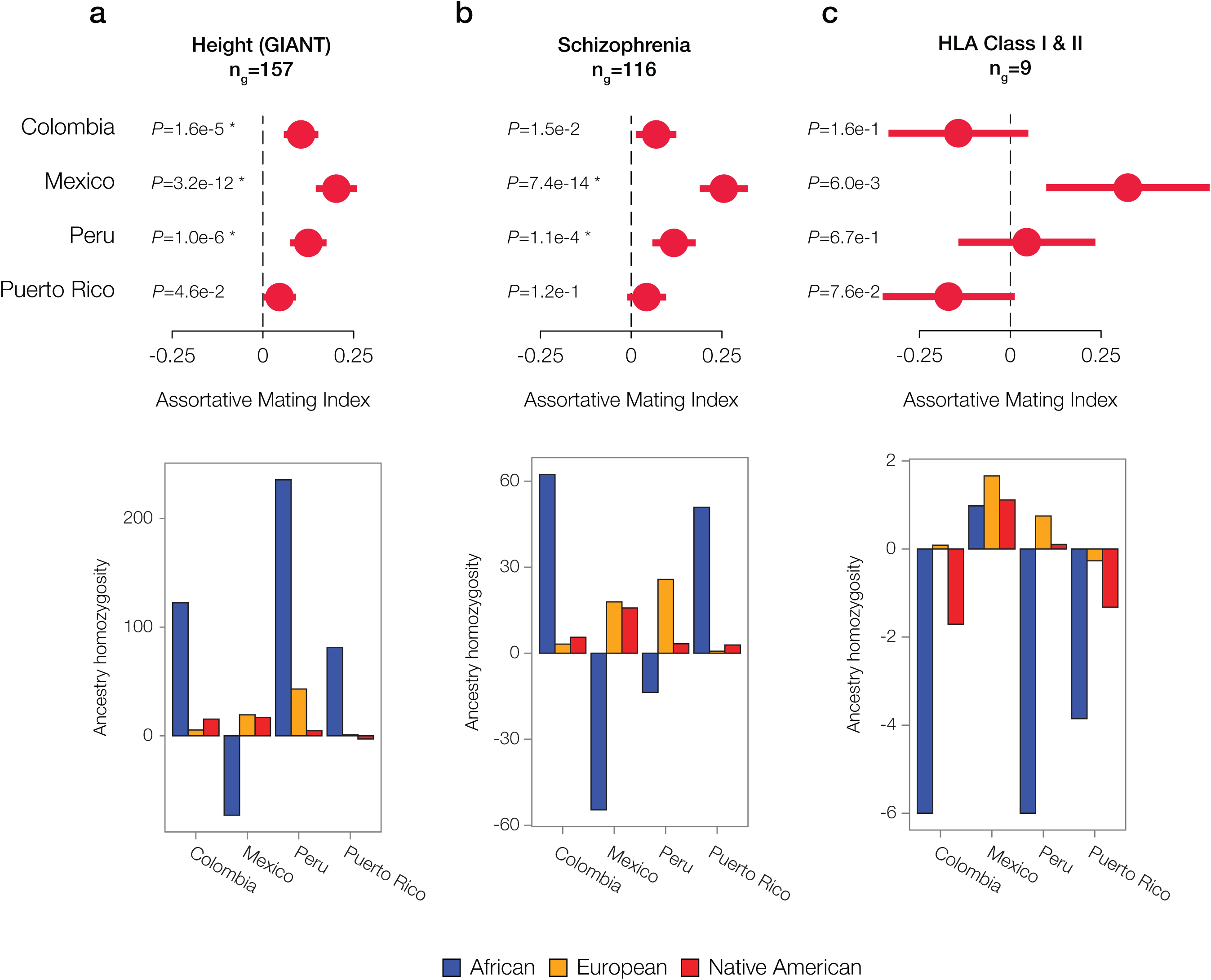
Individual examples of ancestry-based assortative and disassortative mating. Results of meta-analysis of (dis)assortative mating on polygenic phenotypes along with their ancestry drivers are shown for **(a)** an anthropometric trait: height, **(b)** a neurological trait: schizophrenia, and **(c)** the immune-related HLA class I and II genes. The meta-analysis plots show pooled AMI odds ratio values along with their 95% CIs and P-values. Stars indicate false discovery rate q-values<0.05. The ancestry driver plots show the extent to which individual ancestry components – African (blue), European (orange), and Native American (red) – have an excess (>0) or a deficit (<0) of homozygosity.

Both Colombia and Puerto Rico show disassortative mating patterns for all HLA loci (both class I and II genes) (Fig. 5c). The combined AMI values for the HLA loci are only marginally significant but they are among the lowest AMI values seen for any trait evaluated here (Additional file 1: Figure S8), and they are also highly variable among populations (Fig. 4b). HLA loci in Colombia and Puerto Rico show a distinct lack of ancestry homozygosity for almost all ancestry components (Fig. 5c). Mexico and Peru, on the other hand, have some evidence for assortative mating for the HLA loci; Mexico has the highest estimates of ancestry homozygosity at HLA loci for any of the four populations, and Peru has an excess of European and Native American ancestry homozygosity and a deficit of African heterozygosity for these genes. Similar results for two additional anthropometric phenotypes are shown in Additional file 1: Figure S9: body mass index and waist-to-hip ratio adjusted for body mass index. These phenotypes show assortative mating in all four populations, with varying components of ancestral homozygosity driving the relationships. When these results are considered together, African ancestry consistently shows the strongest effect on driving assortative and disassortative mating in admixed Latin American populations (Fig. 5 and Additional file 1: Figure S9).

We further evaluated the extent to which specific ancestry components may drive assortative mating patterns among admixed individuals by evaluating the variance of the three continental ancestry components among individuals within each Latin American population. Assortative mating is known to increase population variance for traits that are involved in mate choice; thus, the ancestry components that drive assortative mating in a given population are expected to show higher overall variance among individual genomes. African ancestry fractions show the highest variation among individuals for all four populations (Fig. 6), consistent with the results seen for the five specific cases of assortative mating evaluated in Fig. 5 and Additional file 1: Figure S9.

**Figure 6.**
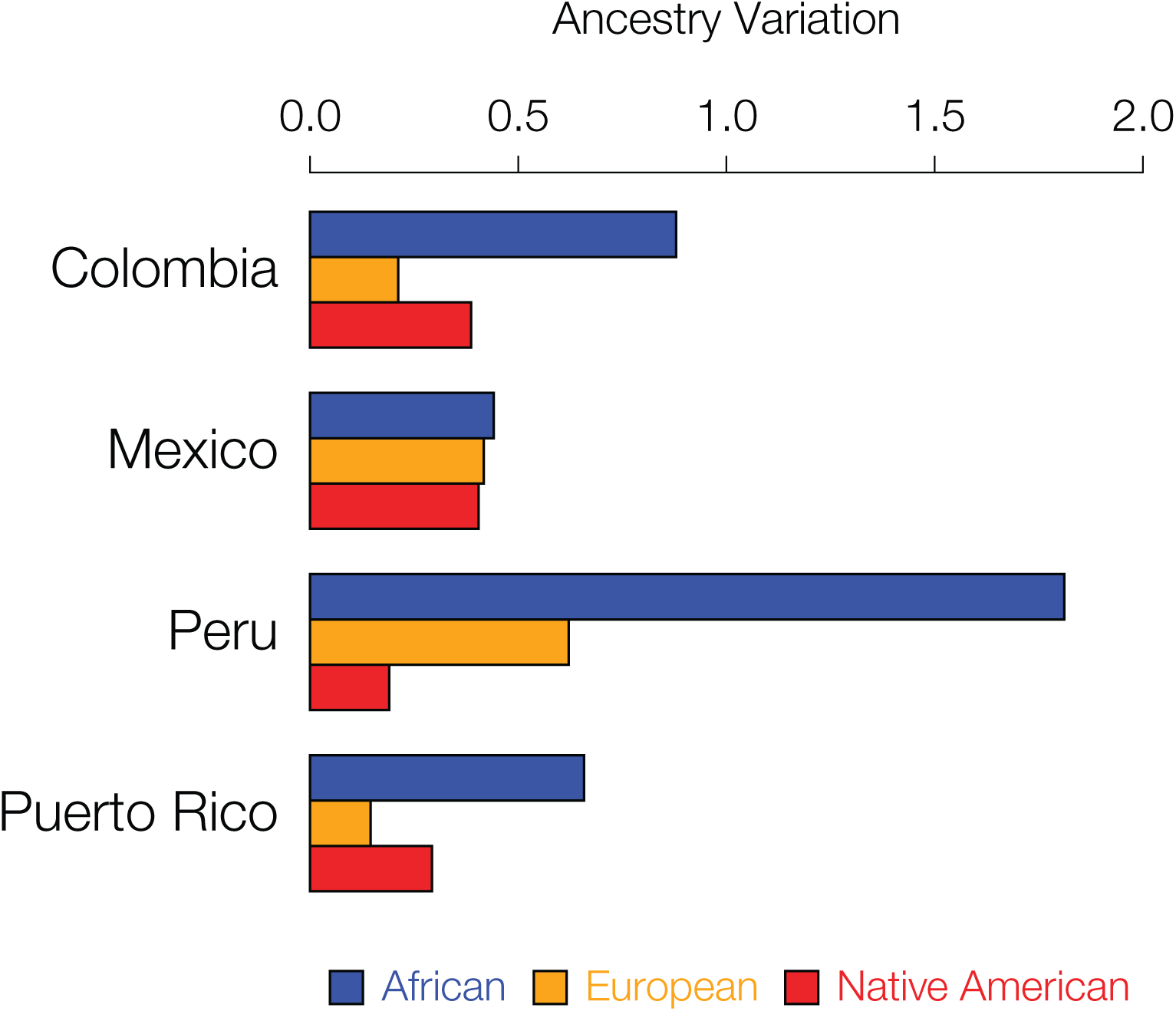
Inter-individual ancestry variance for the four admixed Latin American populations analyzed here. Variance among individuals for the African (blue), European (orange), and Native American (red) ancestry fractions within each population are shown.

## Discussion

Assortative mating is a nearly universal human behavior, and scientists have long been fascinated by the subject [1, 3]. Studies of assortative mating in humans have most often entailed direct measurements of traits – such as physical stature, education and ethnicity – followed by correlation of trait values between partners. Decades of such studies have revealed numerous, widely varying traits that are implicated in mate choice and assortative mating. Studies of this kind typically make no assumptions regarding, nor have any knowledge of, the genetic heredity of the traits under consideration. Moreover, the extent to which the expression of these traits varies among human population groups has largely been ignored.

More recent studies of assortative mating, powered by advances in human genomics, have begun to explore the genetic architecture underlying the human traits that form the basis of mate choice [2, 21]. In addition, recent genomic analyses have underscored the extent to which human genetic ancestry influences assortative mating [24, 30, 31]. However, until this time, these two strands of inquiry have not been brought together. The approach that we developed for this study allowed us to directly assess the connection between local genetic ancestry – *i.e.,* ancestry assignments for specific genome regions or haplotypes – and the human traits that serve as cues for assortative mating.

Our approach relies on the well-established principle that assortative mating results in an excess of genetic homozygosity [29]. However, we do not analyze homozygosity of specific genetic variants *per se*, as is normally done, rather we evaluate excess homozygosity, or the lack thereof, for ancestry-specific haplotypes (Fig. 2b). By merging this approach with data on the genetic architecture of polygenic human phenotypes, we were able to uncover specific traits that inform ancestry-based assortative mating. This is because, when individuals exercise mate choice decisions based on ancestry, they must do so using phenotypic cues that are ancestry-associated. In other words, ancestry-based assortative mating is, by definition, predicated upon traits that vary in expression among human population groups. An obvious example of this is skin color [32], and studies have indeed shown skin color to be an important feature of assortative mating [42-45]. It follows that the assortative mating traits that our study uncovered in admixed Latin American populations must be both genetically heritable and variable among African, European and Native American population groups.

The anthropometric traits found in our study – body mass, height, waist-hip ratio, and facial development – are both heritable and known to vary among the continental population groups that admixed to form modern Latin American populations. This implies that the genetic variants that influence these traits should also vary among these populations. Accordingly, it is readily apparent that mate choice decisions based on these physical features could track local genetic ancestry. Interpretation of the neurological traits that show evidence of local ancestry-based assortative mating – schizophrenia and educational attainment – is not quite as straightforward. For schizophrenia, it is far more likely that we are analyzing genetic loci associated with a spectrum of personality traits that influence assortative mating, as opposed to mate choice based on full-blown schizophrenia, and indeed personality traits are widely known to impact mate choice decisions [19, 22, 25]. In addition, since schizophrenia prevalence does not vary greatly world-wide [46], it is more likely that ancestry-based assortative mating for this trait is tracking an underlying endophenotype rather than the disease itself. While educational attainment outcomes are largely environmentally determined, recent large-scale GWAS studies have uncovered a substantial genetic component to this trait, which is distributed among scores of loci across the genome [47-50]. The population distribution of education associated variants is currently unknown, but our results suggest the possibility of ancestry-variation for some of them.

Mate choice based on divergent MHC loci, apparently driven by body odor preferences, is the best known example of human disassortative mating [28]. However, studies of this phenomenon have largely relied on ethnically homogenous cohorts. In one case where females were asked to select preferred MHC-mediated odors from males of a different ethnic group, they actually preferred odors of males with more similar MHC alleles [51]. Another study showed differences in MHC-dependent mate choice for human populations with distinct ancestry profiles [27]. Ours is the first study that addresses the role of ancestry in MHC-dependent mate choice in ethnically diverse admixed populations. Unexpectedly, we found very different results for MHC-dependent mate choice among the four Latin American populations that we studied. In fact, AMI values for the HLA loci are among the most population variable for any trait analyzed here (Fig. 4b). Mexico and Peru show evidence of assortative mating at HLA loci, whereas Colombia and Puerto Rico show evidence for disassortative mating (Fig. 5c). Interestingly, disassortative mating for HLA loci in Colombia and Puerto Rico is largely driven by African ancestry, and these two populations have substantially higher levels of African ancestry compared to Mexico and Peru. The population- and ancestry-specific dynamics of MHC-dependent mate choice revealed here underscore the complexity of this issue.

Assortative mating alone is not expected to change the frequencies of alleles, or ancestry fractions in the case of our study, within a population. Assortative mating does, however, change genotype frequencies, resulting in an excess of homozygous genotypes. Accordingly, ancestry-based assortative mating is expected to yield an excess of homozygosity for local ancestry assignments (*i.e.*, ancestry-specific haplotypes) (Fig. 2b). By increasing homozygosity in this way, assortative mating also increases the population genetic variance for the traits that influence mate choice. In other words, assortative mating will lead to more extreme, and less intermediate, phenotypes than expected by chance. This population genetic consequence of assortative mating allowed us to evaluate the extent to which specific continental ancestries drive mate choice decisions in admixed populations, since specific ancestry drivers of assortative mating are expected to have increased variance. We found that the fractions of African ancestry have the highest variance among individuals for all four populations, consistent with the idea that traits that are associated with African ancestry drive most of the local ancestry-based assortative mating seen in this study (Fig. 6).

## Conclusions

The confluence of African, European and Native American populations that marked the conquest and colonization of the New World yielded modern Latin American populations that are characterized by three-way genetic admixture [11-15]. Nevertheless, mate choice in Latin America is far from random [24, 31]. Indeed, our results underscore the prevalence of ancestry-based assortative mating in modern Latin American societies. The local ancestry approach that we developed provided new insight into this process by allowing us to hone in on the phenotypic cues that underlie ancestry-based assortative mating. Our method also illuminates the specific ancestry components that drive assortative mating for different traits and makes predictions regarding traits that should vary among continental population groups.

## Methods

### Whole genome sequences and genotypes

Whole genome sequence data for the four admixed Latin American populations studied here were taken from the Phase 3 data release of the 1000 Genomes Project (1KGP) [7]. Whole genome sequence data and genotypes for the putative ancestral populations (Africa, Europe and the Americas) were taken from the 1KGP, the Human Genome Diversity Project [6] (HGDP) and a previous study on Native American genetic ancestry [36].

Whole genome sequence data and genotypes were merged, sites common to all datasets were kept, and single nucleotide polymorphism (SNP) strand orientation was corrected as needed, using PLINK version 1.9 [37]. The resulting dataset consisted of 1,645 individuals from 38 populations with variants characterized for 239,989 SNPs. The set of merged SNP genotypes was phased, using the program SHAPEIT version 2.r837 [38], with the 1KGP haplotype reference panel. This phased set of SNP genotypes was used for local ancestry analysis. PLINK was used to further prune the phased SNPs for linkage, yielding a pruned dataset containing 58,898 linkage-independent SNPs. This pruned set of SNP genotypes was used for global ancestry analysis.

### Global and local ancestry assignment

To infer continental (global) ancestry of the four admixed Latin American populations, ADMIXTURE [39] was run on the pruned SNP genotype dataset (*n*=58,898). ADMIXTURE was run using a K=4, yielding African, European, Asian and Native American ancestry fractions of each admixed individual; the final Asian and Native American fractions were summed to determine the Native American fraction of each individual. For local ancestry analysis of the admixed Latin American populations, the program RFMix [35] version 1.5.4 was run in the PopPhased mode with a minimum node size of 5 and the ‘usereference-panels-in-EM’ option with 2 EM iterations for each individual in the dataset using the phased SNP genotypes (*n*=239,989). Continental African, European, and Native American populations were used as reference populations, and contiguous regions with the same ancestry assignment, *i.e.*, ancestry-specific haplotypes, were delineated where the RFMix ancestry assignment certainty was at least 99%.

Autosomal NCBI RefSeq coding genes were accessed from the UCSC Genome Browser and mapped to the ancestry-specific haplotypes characterized for each admixed Latin American individual. For each diploid genome analyzed here, individual genes can have 0, 1 or 2 ancestry assignments depending on the number of high confidence ancestry-specific haplotypes at that locus. Our assortative mating index (AMI, see below) can only be computed for genes that have 2 ancestry assignments in any given individual, *i.e.,* cases where the ancestry is assigned for both copies of the gene. Thus, for each Latin American population *p*, the mean 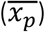 and standard deviation (*sd*_*p*_) of the number of genes with 2 ancestry assignments were calculated and used to compute an ancestry genotype threshold for the inclusion of genes in subsequent analyses. Genes were used in subsequent assortative mating analyses only if they were present above the ancestry genotype threshold of 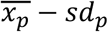.

### Gene sets for polygenic phenotypes

The polygenic genetic architectures of phenotypes that could be effected by assortative mating were characterized using a variety of studies taken from the NHGRI-EBI GWAS Catalog [40], the Genetic Investigation of ANthropometric Traits (GIANT) consortium (http://portals.broadinstitute.org/collaboration/giant/index.php/GIANT_consortium), and PubMed literature sources.

For each polygenic phenotype, all SNPs previously implicated at genome-wide significance levels of *P*≤10^-8^ were collected as the phenotype SNP set. The gene sets for the polygenic phenotypes were collected by directly mapping trait-associated SNPs to genes. SNPs were used to create a gene set only if the SNP fell directly within a gene and thus no intergenic SNPs were used in creating gene sets. Gene sets from the GWAS Catalog were mapped from SNPs using EBI’s in-house pipeline. Sets from GIANT were mapped according to specifications of each individual paper. Gene sets from literature searching were mapped using NCBI’s dbSNP. For each Latin American population, phenotype gene sets were filtered to only include genes that passed the ancestry genotype threshold, as described previously. Finally, the polygenic phenotype gene sets were filtered based on size, so that all polygenic phenotypes included two or more genes. The final data set contains gene sets for 106 polygenic phenotypes, hierarchically organized into three functional categories, including 986 unique genes (Additional file 1: Figure S5).

### Assortative mating index (AMI)

To assess local ancestry-based assortative mating, we developed the assortative mating index (AMI), a log odds ratio test statistic that computes the relative local ancestry homozygosity compared to heterozygosity for any given gene. Ancestry homozygosity occurs when both genes in a genome have the same local ancestry, whereas ancestry heterozygosity refers to a pair of genes in a genome with different local ancestry assignments. The assortative mating index (*AMI*) is calculated as:

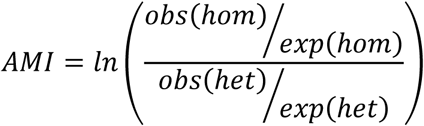

where ^*obs*(*hom*)^/_*exp* (*hom*)_ is the ratio of the observed and expected local ancestry homozygous gene pairs and ^*obs*(*het*)^/_*exp* (*het*)_ is the ratio of the observed and expected local ancestry heterozygous gene pairs.

The observed values of local ancestry homozygous and heterozygous gene pairs are taken from the gene-to-ancestry mapping data for each gene in each population. The expected values of local ancestry homozygous and heterozygous gene pairs are calculated for each gene in a population using a triallelic Hardy-Weinberg (HW) model, in which the gene-specific local ancestry assignment fractions are taken as the three allele frequencies. For the African (*a*), European (*e*), and Native American (*n*) gene-specific local ancestry assignment fractions in a population, the HW expected genotype frequencies are: (*a* + *e* + *n*)^2^ or *a*^2^ + 2*ae* + *e*^2^ + 2*an* + 2*en* + *n*^2^. Accordingly, the expected frequency of homozygous pairs is *a*^2^ +*e*^2^ + *n*^2^ and the expected frequency of heterozygous pairs is 2*ae* + 2*an* + 2*en*. For each gene, in each population, the expected homozygous and heterozygous frequencies are multiplied by the number of individuals with two ancestry assignments for that gene to yield the expected counts of gene pairs in each class.

For each polygenic phenotype, a meta-analysis of gene-specific AMI values was conducted to evaluate the effect of all of the genes involved in the phenotype on assortative mating, using the metafor [41] package in R. 95% confidence intervals for each gene, meta-gene AMI values, significance *P*-values, and false discovery rate *q*-values, were computed using the Mantel-Haenszel method under a fixed-effects model.

### Permutation of random mating

A standard permutation testing framework was adopted for the approximation of random mating in each of the four Latin American populations. Random mating was approximated by randomly combining pairs of individual phased haplotypes from a population to yield permuted diploid genotypes. Haploid chromosomes were permuted randomly within each population using the Fisher-Yates shuffle. After permutation of the chromosomes, per gene AMI values were re-calculated for all genes passing the population-specific ancestry genotyping thresholds. The permutations were completed 20 times, and the population-specific mean AMI values for each gene were taken as the permuted AMI for the gene. This mean permuted AMI per gene was used in AMI meta-analysis for each gene set to determine expected AMI values.

### Population genetic simulation of assortative mating

To validate the performance of the AMI test statistic, we adopted a population genetic model that simulates assortative mating in the four Latin American populations under Hardy-Weinberg equilibrium, with a fraction of the population mating assortatively. For each gene in a given population, the present-day local ancestry assignment fractions are used as the starting ancestral proportions: African = *a*, European = *e*, Native American = *n*. Using a triallelic Hardy-Weinberg model, taking the ancestral proportions as the allele frequencies, the ancestry genotype frequencies for a given gene at the starting generation are calculated as:

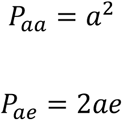

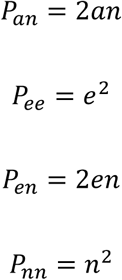

where *P*_*aa*_ = African-African genotype, *P*_*ae*_ = African-European genotype, *P*_*an*_ = African-Native American genotype, *P*_*ee*_ = European-European genotype, *P*_*en*_ = European-Native American genotype and *P*_*nn*_ = Native American-Native American genotype. Under the model, the fraction of the population that mates assortatively is denoted as α and the fraction that mates randomly is 1 - α. Taking the current generation ancestry genotype frequencies, the subsequent generation’s ancestry genotype frequencies are calculated using the formulae:

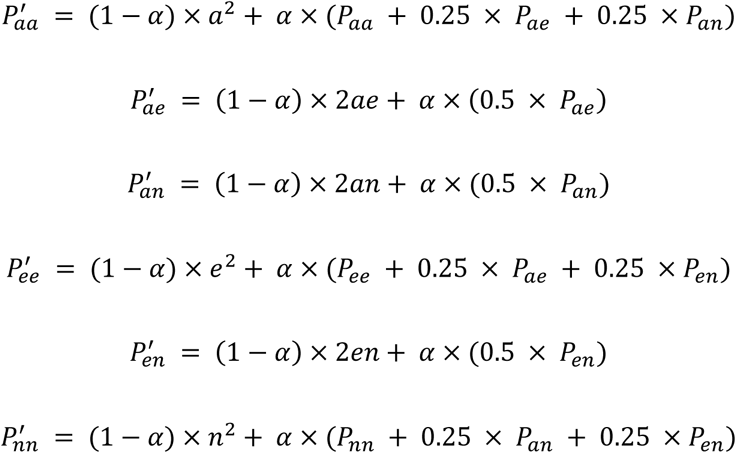

Ancestry genotypes in each population were simulated for 20 generations, with the assumption of a generation time of 25 years, accounting for 500 years of elapsed time during the conquest and colonization of the Americas. The final ancestry genotype frequencies after the 20 generations were used to calculate the simulated ancestry homozygosity and heterozygosity values. For each Latin American simulated population, random gene sets, ranging in size from 2 to 20, were created by subsampling genes in the simulation. A meta-analysis AMI value and *P*-value for each gene set was calculated using the fixed-effects model of the Mantel-Haenszel method.

### Ancestry-specific drivers of assortative mating

For each significant polygenic phenotype of interest, we identified the ancestry component related to mate choice by calculating the ancestry homozygosity 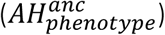 for all genes for each ancestry at the given phenotype. The ancestry homozygosity was calculated as

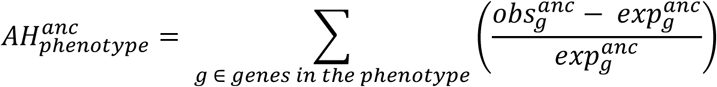

where is *anc* one of the three ancestries – African, European or Native American, *g* ∈ *genes in the phenotype* are all of the genes involved in the polygenic *phenotype*, 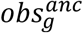 is the number of observed homozygous genes for gene *g* coming from *anc*, and 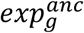 is the number of expected homozygous genes for gene *g* coming from *anc* (as calculated using a triallelic Hardy-Weinberg model).

### Statistical significance testing

Significance testing for the difference between the observed and expected AMI distributions was completed using the t-test package in R. The metafor package, used for calculating the meta-analysis AMI values, also calculates a *P*-value and a false discovery rate *q*-value to correct for multiple statistical tests, which were used for identifying polygenic phenotypes that are significantly influenced by local ancestry-based assortative mating in each Latin American population. The variance of AMI values across the four populations for each phenotype was calculated as it is implemented in R and used for identifying phenotypes that had highly similar (minimal variance) or highly dissimilar (maximal variance) local ancestry-based assortative mating patterns. The coefficient of variation was used to measure the inter-individual variance for each of the three continental ancestry components within the four admixed Latin American populations analyzed here.

## Declarations

### Ethics approval and consent to participate

The de-identified human genome sequence data analyzed here are made publicly available as part of the 1000 Genomes Project and the Human Genome Diversity Project.

### Consent for publication

Not applicable

### Availability of data and materials

1000 Genomes Project data are available from http://www.internationalgenome.org/data/

Human Genome Diversity Project data are available from http://www.hagsc.org/hgdp/

Previously published Native American genotype data can be accessed from a data use agreement governed by the University of Antioquia as previously described [36].

### Authors’ contributions

ETN conducted all of the ancestry-based assortative mating analyses. LR performed the permutation and simulation analyses. LW participated in the assortative mating analysis for individual phenotypes. ABC performed the genetic ancestry analyses. ETN, ATC, AMD and AV curated the GWAS SNP associations and polygenic phenotype gene sets. ETN, LR, LW and ABC generated the manuscript figures. IKJ conceived of, designed and supervised the project. ETN, LR and IKJ wrote the manuscript. All authors read and approved the final manuscript.

## Additional files

**Additional file 1: Figure S1.** Global locations of the populations analyzed in this study. **Figure S2.** Three-way continental genetic ancestry for the four admixed Latin American populations analyzed in this study. **Figure S3.** Local ancestry assignment with chromosome painting. **Figure S4.** Comparison of ancestry fractions estimated by ADMIXTURE (global ancestry) versus RFMix (local ancestry). **Figure S5.** Polygenic phenotypes taken from genome-wide association studies (GWAS). **Figure S6.** Distributions of observed (dark blue) versus expected (light blue) AMI values for the four admixed Latin American populations analyzed here. **Figure S7.** Simulation of the assortative mating index (AMI) test statistic under assortative mating. **Figure S8.** Assortative mating index (AMI) values for all phenotypes across all four populations analyzed here. **Figure S9.** Individual examples of ancestry-based assortative mating. **Table S1**. References and values for phenotypes with significant AMI values and population variance.

## References

1. Vandenburg SG: Assortative mating, or who marries whom? Behav Genet 1972, 2:127–157.

2. Robinson MR, Kleinman A, Graff M, Vinkhuyzen AA, Couper D, Miller MB, Peyrot WJ, Abdellaoui A, Zietsch BP, Nolte IM: Genetic evidence of assortative mating in humans. Nature Human Behaviour 2017, 1:0016.

3. Buss DM: Human mate selection: Opposites are sometimes said to attract, but in fact we are likely to marry someone who is similar to us in almost every variable. American scientist 1985, 73:47–51.

4. Cavalli-Sforza LL, Menozzi P, Piazza A: The history and geography of human genes. Princeton: Princeton University Press; 1994.

5. Rosenberg NA, Pritchard JK, Weber JL, Cann HM, Kidd KK, Zhivotovsky LA, Feldman MW: Genetic structure of human populations. Science 2002, 298:2381–2385.

6. Li JZ, Absher DM, Tang H, Southwick AM, Casto AM, Ramachandran S, Cann HM, Barsh GS, Feldman M, Cavalli-Sforza LL, Myers RM: Worldwide human relationships inferred from genome-wide patterns of variation. Science 2008, 319:1100–1104.

7. Genomes Project C, Auton A, Brooks LD, Durbin RM, Garrison EP, Kang HM, Korbel JO, Marchini JL, McCarthy S, McVean GA, Abecasis GR: A global reference for human genetic variation. Nature 2015, 526:68–74.

8. Hellenthal G, Busby GB, Band G, Wilson JF, Capelli C, Falush D, Myers S: A genetic atlas of human admixture history. Science 2014, 343:747–751.

9. Mann CC: 1493: Uncovering the new world Columbus created. New York: Vintage; 2011.

10. Jordan IK: The Columbian Exchange as a source of adaptive introgression in human populations. Biol Direct 2016, 11:17.

11. Montinaro F, Busby GB, Pascali VL, Myers S, Hellenthal G, Capelli C: Unravelling the hidden ancestry of American admixed populations. Nat Commun 2015, 6:6596.

12. Rishishwar L, Conley AB, Wigington CH, Wang L, Valderrama-Aguirre A, Jordan IK: Ancestry, admixture and fitness in Colombian genomes. Sci Rep 2015, 5:12376.

13. Wang S, Ray N, Rojas W, Parra MV, Bedoya G, Gallo C, Poletti G, Mazzotti G, Hill K, Hurtado AM, et al: Geographic patterns of genome admixture in Latin American Mestizos. PLoS Genet 2008, 4:e1000037.

14. Ruiz-Linares A, Adhikari K, Acuna-Alonzo V, Quinto-Sanchez M, Jaramillo C, Arias W, Fuentes M, Pizarro M, Everardo P, de Avila F, et al: Admixture in Latin America: geographic structure, phenotypic diversity and self-perception of ancestry based on 7,342 individuals. PLoS Genet 2014, 10:e1004572.

15. Bryc K, Velez C, Karafet T, Moreno-Estrada A, Reynolds A, Auton A, Hammer M, Bustamante CD, Ostrer H: Colloquium paper: genome-wide patterns of population structure and admixture among Hispanic/Latino populations. Proc Natl Acad Sci U S A 2010, 107 Suppl 2:8954–8961.

16. Allison DB, Neale MC, Kezis MI, Alfonso VC, Heshka S, Heymsfield SB: Assortative mating for relative weight: genetic implications. Behav Genet 1996, 26:103–111.

17. Silventoinen K, Kaprio J, Lahelma E, Viken RJ, Rose RJ: Assortative mating by body height and BMI: Finnish twins and their spouses. Am J Hum Biol 2003, 15:620–627.

18. Stulp G, Simons MJ, Grasman S, Pollet TV: Assortative mating for human height: A meta-analysis. Am J Hum Biol 2017, 29.

19. Merikangas KR: Assortative mating for psychiatric disorders and psychological traits. Arch Gen Psychiatry 1982, 39:1173–1180.

20. Mare RD: Five decades of educational assortative mating. American sociological review 1991:15–32.

21. Domingue BW, Fletcher J, Conley D, Boardman JD: Genetic and educational assortative mating among US adults. Proc Natl Acad Sci U S A 2014, 111:7996–8000.

22. Hur YM: Assortive mating for personaltiy traits, educational level, religious affiliation, height, weight, adn body mass index in parents of Korean twin sample. Twin Res 2003, 6:467–470.

23. Salces I, Rebato E, Susanne C: Evidence of phenotypic and social assortative mating for anthropometric and physiological traits in couples from the Basque country (Spain). J Biosoc Sci 2004, 36:235–250.

24. Zou JY, Park DS, Burchard EG, Torgerson DG, Pino-Yanes M, Song YS, Sankararaman S, Halperin E, Zaitlen N: Genetic and socioeconomic study of mate choice in Latinos reveals novel assortment patterns. Proc Natl Acad Sci U S A 2015, 112:13621–13626.

25. Kandler C, Bleidorn W, Riemann R: Left or right? Sources of political orientation: the roles of genetic factors, cultural transmission, assortative mating, and personality. J Pers Soc Psychol 2012, 102:633–645.

26. Kalmijn M: Assortative mating by cultural and economic occupational status. American Journal of Sociology 1994, 100:422–452.

27. Chaix R, Cao C, Donnelly P: Is mate choice in humans MHC-dependent? PLoS Genet 2008, 4:e1000184.

28. Wedekind C, Seebeck T, Bettens F, Paepke AJ: MHC-dependent mate preferences in humans. Proc Biol Sci 1995, 260:245–249.

29. Sebro R, Hoffman TJ, Lange C, Rogus JJ, Risch NJ: Testing for non-random mating: evidence for ancestry-related assortative mating in the Framingham heart study. Genet Epidemiol 2010, 34:674–679.

30. Zaitlen N, Huntsman S, Hu D, Spear M, Eng C, Oh SS, White MJ, Mak A, Davis A, Meade K, et al: The Effects of Migration and Assortative Mating on Admixture Linkage Disequilibrium. Genetics 2017, 205:375–383.

31. Risch N, Choudhry S, Via M, Basu A, Sebro R, Eng C, Beckman K, Thyne S, Chapela R, Rodriguez-Santana JR, et al: Ancestry-related assortative mating in Latino populations. Genome Biol 2009, 10:R132.

32. Hancock AM, Witonsky DB, Alkorta-Aranburu G, Beall CM, Gebremedhin A, Sukernik R, Utermann G, Pritchard JK, Coop G, Di Rienzo A: Adaptations to climate-mediated selective pressures in humans. PLoS Genet 2011, 7:e1001375.

33. Homburger JR, Moreno-Estrada A, Gignoux CR, Nelson D, Sanchez E, Ortiz-Tello P, Pons-Estel BA, Acevedo-Vasquez E, Miranda P, Langefeld CD, et al: Genomic Insights into the Ancestry and Demographic History of South America. PLoS Genet 2015, 11:e1005602.

34. Moreno-Estrada A, Gravel S, Zakharia F, McCauley JL, Byrnes JK, Gignoux CR, Ortiz-Tello PA, Martinez RJ, Hedges DJ, Morris RW, et al: Reconstructing the population genetic history of the Caribbean. PLoS Genet 2013, 9:e1003925.

35. Maples BK, Gravel S, Kenny EE, Bustamante CD: RFMix: a discriminative modeling approach for rapid and robust local-ancestry inference. Am J Hum Genet 2013, 93:278–288.

36. Reich D, Patterson N, Campbell D, Tandon A, Mazieres S, Ray N, Parra MV, Rojas W, Duque C, Mesa N, et al: Reconstructing Native American population history. Nature 2012, 488:370–374.

37. Chang CC, Chow CC, Tellier LC, Vattikuti S, Purcell SM, Lee JJ: Second-generation PLINK: rising to the challenge of larger and richer datasets. Gigascience 2015, 4:7.

38. Delaneau O, Zagury JF, Marchini J: Improved whole-chromosome phasing for disease and population genetic studies. Nat Methods 2013, 10:5–6.

39. Alexander DH, Novembre J, Lange K: Fast model-based estimation of ancestry in unrelated individuals. Genome Res 2009, 19:1655–1664.

40. MacArthur J, Bowler E, Cerezo M, Gil L, Hall P, Hastings E, Junkins H, McMahon A, Milano A, Morales J, et al: The new NHGRI-EBI Catalog of published genome-wide association studies (GWAS Catalog). Nucleic Acids Res 2017, 45:D896–D901.

41. Viechtbauer W: Conducting meta-analyses in R with the metafor package. Journal of Statistical Software 2010, 36:1–48.

42. Procidano ME, Rogler LH: Homogamous assortative mating among Puerto Rican families: intergenerational processes and the migration experience. Behav Genet 1989, 19:343–354.

43. Trachtenberg A, Stark AE, Salzano FM, Da Rocha FJ: Canonical correlation analysis of assortative mating in two groups of Brazilians. J Biosoc Sci 1985, 17:389–403.

44. Malina RM, Selby HA, Buschang PH, Aronson WL, Little BB: Assortative mating for phenotypic characteristics in a Zapotec community in Oaxaca, Mexico. J Biosoc Sci 1983, 15:273–280.

45. Frisancho AR, Wainwright R, Way A: Heritability and components of phenotypic expression in skin reflectance of Mestizos from the Peruvian lowlands. Am J Phys Anthropol 1981, 55:203–208.

46. Saha S, Chant D, Welham J, McGrath J: A systematic review of the prevalence of schizophrenia. PLoS Med 2005, 2:e141.

47. Davies G, Marioni RE, Liewald DC, Hill WD, Hagenaars SP, Harris SE, Ritchie SJ, Luciano M, Fawns-Ritchie C, Lyall D, et al: Genome-wide association study of cognitive functions and educational attainment in UK Biobank (N=112 151). Mol Psychiatry 2016, 21:758–767.

48. Okbay A, Beauchamp JP, Fontana MA, Lee JJ, Pers TH, Rietveld CA, Turley P, Chen GB, Emilsson V, Meddens SF, et al: Genome-wide association study identifies 74 loci associated with educational attainment. Nature 2016, 533:539–542.

49. Rietveld CA, Esko T, Davies G, Pers TH, Turley P, Benyamin B, Chabris CF, Emilsson V, Johnson AD, Lee JJ, et al: Common genetic variants associated with cognitive performance identified using the proxy-phenotype method. Proc Natl Acad Sci U S A 2014, 111:13790–13794.

50. Rietveld CA, Medland SE, Derringer J, Yang J, Esko T, Martin NW, Westra HJ, Shakhbazov K, Abdellaoui A, Agrawal A, et al: GWAS of 126,559 individuals identifies genetic variants associated with educational attainment. Science 2013, 340:1467–1471.

51. Jacob S, McClintock MK, Zelano B, Ober C: Paternally inherited HLA alleles are associated with women’s choice of male odor. Nat Genet 2002, 30:175–179.

